# Asparagine endopeptidase cleaves tau at N167 after uptake into microglia

**DOI:** 10.1101/560110

**Authors:** Annika Behrendt, Maria Bichmann, Ebru Ercan-Herbst, Per Haberkant, David C. Schöndorf, Michael Wolf, Salma A. Fahim, Enrico Murolo, Dagmar E. Ehrnhoefer

## Abstract

**Background:** Tau cleavage by different proteolytic enzymes generates short, aggregation-prone fragments that have been implicated in the pathogenesis of Alzheimer’s disease (AD). Asparagine endopeptidase (AEP) activity in particular has been associated with tau dysfunction and aggregation, and the activity of the protease is increased in both aging and AD.

**Methods and Results:** Using a mass spectrometry approach we identified a novel tau cleavage site at N167 and confirmed its processing by AEP. In combination with the previously known site at N368, we show that AEP cleavage yields a tau fragment that is present in both control and AD brains at similar levels. AEP is a lysosomal enzyme, and our data suggest that it is expressed in microglia rather than in neurons. Accordingly, we observe tau cleavage at N167 and N368 after endocytotic uptake into microglia, but not neurons. However, tau_168-368_ does not accumulate in microglia and we thus conclude that the fragment is part of a proteolytic cascade leading to tau degradation.

**Conclusions:** While we confirm previous studies showing increased overall AEP activity in AD brains, our data suggests that AEP-mediated cleavage of tau is a physiological event occurring during microglial degradation of the secreted neuronal protein. The disease-associated increase in active AEP may thus be related to pro-inflammatory conditions in AD brains, and our findings argue against AEP inhibition as a therapeutic approach in AD.

## Background

Intraneuronal aggregates consisting of the microtubule-associated protein tau are associated with several neurodegenerative diseases, including Alzheimer’s disease (AD). The aggregation process is promoted by several post-translational modifications, including proteolytic events leading to short tau fragments that are more prone to misfolding (1, 2). In AD, tau pathology spreads throughout the brain in a stereotypical manner along synaptically connected brain areas (3). However, the process of tau secretion and uptake also occurs physiologically and can be regulated by neuronal activity (4, 5). Multiple pathways for tau secretion have been described, including exosomal and non-conventional secretion of free protein (6–8), while uptake is thought to occur mainly through endocytosis (4, 9, 10).

However, neuropathological studies have shown that the endosomal-lysosomal pathway is dysfunctional in AD, and these disturbances might play a role for both amyloid and tau processing (11). Among the lysosomal enzymes, asparagine endopeptidase (AEP) was shown to cleave tau at N368, and the cleavage product was identified by immunostaining as well as mass spectrometry in human brain (12). Cleavage was also shown in the P301S mouse model of tauopathy and was blocked upon genetic ablation of AEP (12). AEP is a lysosomal enzyme that requires a low pH for activation (13). Although cytoplasmic activity has been demonstrated under certain conditions (14), it is mostly described for its role in the endolysosomal system of dendritic cells (15). The activity of AEP in the brain increases with age and is also elevated in AD, leading to the cleavage of additional disease proteins such as amyloid precursor protein (APP) or α-synuclein (12, 16). AEP has therefore been proposed as a potential therapeutic target for multiple neurodegenerative diseases (17).

While tau processing, spreading and uptake is mostly studied in neurons, other cell types present in the brain might contribute. In particular, microglia as the resident phagocytes in the brain are a major player in the clearance of protein aggregates, dysfunctional neurons and the pruning of synapses in neurodegeneration (18). Accordingly, microglia have been shown to take up recombinant as well as human brain-derived tau aggregates in vitro and in vivo in mouse models (19), as well as phagocytose whole neurons containing tau aggregates (20). Microglia may furthermore contribute to the propagation of tau pathology via exosome secretion in vivo, since the depletion of microglia and inhibition of exosome synthesis decreases tau spreading in mice (21). However, it is not yet fully elucidated if microglia further process tau for example by cleavage, which may generate fragments prone to misfolding and aggregation. The finding that tau seeds derived from human AD patients or Tg4510 mice model are broken down in microglia (22) nevertheless indicates that processing in the endosomal-lysosomal compartment can occur.

In this study we used a mass spectrometry approach to determine tau cleavage sites in human brain, using both control and AD samples. We identified a novel proteolytic event at N167 and demonstrate that AEP is able to process tau at this site in addition to the previously published cleavage at N368. We find that tau is processed in a pH-dependent manner upon endocytosis, and show that microglia, rather than neurons, possess high levels of AEP and related tau cleavage activity. While we confirm previous reports of increased AEP activity in AD patient brain tissue, our results point towards a role for AEP in the physiological degradation of tau secreted by neurons.

## Methods

### Recombinant human tau purification

All tau variants (full length protein and fragment encoding amino acids 168-368 or amino acids 256-368) were cloned into the pET19b vector (Novagen) in between the NcoI and BamHI restriction sites. The pET19b-Tau plasmids were transformed into E. coli BL21(DE3) cells (Novagen). Cells were grown in LB supplemented with ampicillin at 37°C until OD600 ∼ 0.6-0.8. The expression of the Tau proteins was induced by the addition of 1 mM IPTG. The cells were then grown for an additional 3 h at 37°C and harvested by centrifugation. The cell pellet was resuspended in running buffer (50 mM Na-phosphate pH 7.0, 1 mM EGTA and 1 mM DTT) supplemented with cOmplete protease inhibitors (Roche), benzonase (Merck) and 10 µg/ml lysozyme (Sigma). The cells were lysed by 4 passages through an EmulsiFlex C3 (Avestin). After centrifugation and filtration, the cleared lysates were boiled for 20 min at 100°C. After another centrifugation and filtration step the lysate was then loaded onto a combination of a HiTrap Q and a HiTrap SP column (GE Healthcare) pre-equilibrated with running buffer. After loading the sample, the HiTrap Q column was removed. The HiTrap SP column was washed with running buffer and eluted in a gradient to running buffer containing 300 mM NaCl. The HiTrap SP elution fractions containing the Tau proteins were concentrated using a 30 MWCO (for full length tau) or 3 MWCO (for tau fragments) Amicon centrifugal filter unit (Merck) and loaded on a HiLoad 16/600 Superdex 75 pg size exclusion chromatography column (GE Healthcare) equilibrated with running buffer. After SDS-PAGE analysis, the elution fractions containing the purest Tau proteins were pooled and quantified. The samples were flash-frozen in liquid nitrogen and the aliquots were stored at −80°C.

### Thioflavin T aggregation assay

10 µM recombinant tau protein were incubated at 37°C in a black 96 well plate in 20 mM Tris pH 7.5 containing 100 mM NaCl, 1 mM EDTA, 1 mM DTT, 0.03 mg/mL heparin sodium salt (Sigma Aldrich) and 30 µM thioflavin T (Sigma Aldrich). Signals were measured every 30 minutes for a time frame of 62 hours using FLUOstar optima (BMG Labtech) (450_Ex_/520_Em_).

Tau_168-368_ seeds were prepared from a 25 µM aggregation reaction, which was stopped well after reaching the signal plateau (24 to 30 h) by shock-freezing in liquid nitrogen; aggregates were then stored at −80°C. To prepare smaller seeds, tau_168-368_ aggregates were sonicated for 10 sec in a Sonorex super RK 106 Ultrasonic bath (Bandelin) and 0.5 µM seeds were added to aggregation reactions with tau_1-441_.

### Biotinylation of recombinant tau

10 mM Biotin solution was prepared from EZ-Link Sulfo NHS-Biotin (Thermo Fisher). The solution was added to a 20-fold molar excess to 72 µM recombinant tau_1-441_ and incubated for 2 h on ice. The excess of non-reacted biotin was removed via Zeba Spin Desalting Columns (Thermo Fisher). To this end, the empty column was washed three times with PBS and centrifuged for 1 min at 1500x g. The biotinylated tau was added and centrifuged for 2 min at 1500x g. The flow through was collected and stored at −20°C until further use.

### Cell culture

HEK T293 cells were cultured at 37°C in 5% CO_2_ in DMEM+GlutaMax (Thermo Fisher) with 10% fetal bovine serum (Sigma) and 1% penicillin/streptomycin (Thermo Fisher). Cells were used for transfections between passage 10-20. Transfection was performed at a cell confluency of 60-70% using JetPrime reagent (PolyPlus) according to manufacturer’s instruction. For each construct, 1 µg DNA (wt 2N4R tau, N167Q 2N4R tau, N368Q 2N4R tau or AEP (Origene clone no. RC200309)) was used. Tau constructs contained an N-terminal HA tag and were cloned into the pcDNA3 vector, mutants were generated by site-directed mutagenesis and verified by sequencing. Cells were transfected and incubated for 20 h before treatment with the lysosomal acidification inhibitor NH_4_Cl (20 mM) or mock-treatment with equal amounts of PBS for additional 6 h. Cells were harvested by scraping, centrifuged for 5 min at 2000x g at 4°C, washed once with ice-cold 1X PBS and centrifuged for 5 min at 2000x g at 4°C. Samples were stored at −20°C until further use.

Cells were lysed in Triton buffer (150 mM NaCl; 20 mM Tris, pH 7.5; 1 mM EDTA; 1 mM EGTA; 1 % Triton-X-100; 1X cOmplete Protease inhibitors; 1X PhosStop Phosphatase inhibitors (Roche)). Cell lysis was promoted by scraping the tubes over a hard surface and incubating the samples on ice for 10 min for three cycles. Samples were centrifuged for 15 min at 12.000x g at 4°C to remove cellular debris and the supernatant was transferred to a fresh Eppendorf tube. The protein concentration was measured using the BCA protein assay according to manufacturer’s instruction (BioRad).

### Immunoprecipitation of microglia media

20 µl magnetic Protein G beads per sample (Invitrogen) were washed two times with 50% BSA-free blocking buffer (Pierce), then incubated with 1 µg Tau5 antibody (Abcam, cat no ab80579) in 200 µl 50% BSA-free blocking buffer for 30 min at room temperature under constant rotation. Coated beads were washed with microglia medium and incubated with 250 µl conditioned media over night at 4°C under constant rotation. Beads were then washed twice with PBS-T, suspended in 10 µl Lämmli sample buffer and boiled for 10 min at 95°C before separation by SDS-PAGE.

### Immunoblotting

Lysates were prepared by boiling at 95°C for 10 minutes in Lämmli sample buffer and separated by SDS gel electrophoresis, followed by immunoblotting on PVDF membranes (Merck Millipore). The membranes were blocked in Odyssey blocking buffer (TBS) (Li-Cor Biosciences) and incubated with primary antibodies overnight at 4°C: Tau5, mouse, 1:1000, cat no ab80579 (Abcam); Tau N368, rabbit, 1:5000, cat no ABN1703 (Merck); HA, mouse, 1:1000, cat no ab18181 (Abcam); DAKO-Tau, rabbit, 1:10000, cat no A0024 (Dako/Agilent); Tau-CS, rabbit, 1:1000, cat no 46687 (Cell Signaling Technology); AEP, goat, 1:200, cat no PA5-47271 (Thermo Fisher); Actin, mouse, 1:5000, cat no A5441 (Sigma Aldrich); Tau12, mouse, 1:500, cat no 806501 (Biolegend); GAPDH, 1:5000, cat no 2118 (Cell Signaling Technology). Antibodies were diluted in 5% BSA in 1X TBST (1X TBS, 0.05% Tween-20). The next day, membranes were washed three times with 1X TBST for 10 min each. Afterwards, secondary antibodies (all 1:20.000, IRDye Donkey anti-mouse 800, IRDye Donkey anti-rabbit 680, IRDye Donkey anti-goat 680) or IRDye Streptavidin 800CW (all Li-cor Biosciences) were diluted in 5% BSA in 1X TBS and incubated for 1 h at room temperature. The membranes were then washed three times with 1X TBST and once with 1X TBS and developed on a Li-Cor Odyssey CLx imaging station (Li-cor Biosciences).

### Neuron culture

Ngn2 induced neurons were differentiated as previously described with minor modifications (23). A doxycycline inducible NGN2 expression cassette was stable integrated in the AAV1 locus using TALEN technology by Bioneer (Denmark). iPSCs were split in a concentration of 100 000 cells/cm^2^ on matrigel (BD) coated plates in mTesR (Stem Cell Technologies). At day 1 after splitting, the medium was changed to N2/B27 medium (50% DMEM/F12, 50% Neurobasal, 1:200 N2, 1:100 B27, 1% PenStrep, 0.5 mM Non-essential amino acids, (all Invitrogen), 50 µM ß-mercaptoethanol (Gibco), 2.5 µg/ml insulin and 1 mM sodium pyruvate (both Sigma)) with 2 µg/ml doxycycline (Sigma). The medium was changed daily. On day 4, cells were split with accutase (Invitrogen) and re-seeded in a density of 200 000 cells/cm^2^ in N2/B27 medium with doxycycline and 10 µM Rock inhibitor Y-27632 (Selleckchem) on matrigel coated plates in the final format. N2/B27 with doxycycline was changed daily until day 7. On day 8 the medium was switched to final maturation medium (FMM; N2B27 with 20 ng/ml BDNF, 10 ng/ml GDNF (both Peprotech), 1 mM dibutyryl-cAMP (Sigma) and 200 µM ascorbic acid (Sigma)). The medium was changed every third day until cells were used for analysis at day 21. To study tau uptake within neurons, biotinylated recombinant tau_1-441_ (3 µM) was added to the cell culture media for 6 hours. Neurons were then harvested and lysates analyzed for recombinant tau uptake and possible tau cleavage.

For experiments examining tau release and uptake in microglia, media from neurons was collected on day 15 of the differentiation, centrifuged at 2000x g for 5 minutes at 4°C and supernatants were stored at −80°C until further use.

### Microglia culture

Immortalized mouse microglia were obtained from Merck Millipore (SCC134) and cultured according to manufacturer’s instructions. To study tau uptake, 800 000 cells were seeded in a 12 well plate and recombinant tau_1-441_ (0.5 µM) was added to the cell media for 6 or 24 h. NH_4_Cl was added to a final concentration of 20 mM where indicated. Cells were harvested and lysed as described above. Conditioned media were collected, centrifuged at 2000x g for 5 min at 4°C and stored at −80°C until use. 250 µl media were subjected to immunoprecipitation with Tau5 antibody as described above.

For experiments with tau released from iPS-derived neurons, microglia media were removed completely and replaced by conditioned neuronal media or fresh FMM as a control. Microglia were incubated with the neuronal media for 24 h, and 20 mM NH_4_Cl were added in the last 6 h where indicated. Conditioned media were collected, centrifuged at 2000x g for 5 min at 4°C and stored at −80°C until use.

### Electrochemiluminescence ELISA

To measure the tau levels in neuronal and microglial media, the V-Plex Human Total Tau Kit (Meso Scale Discovery, Cat. No: K151LAE) was used, following the manufacturer’s protocol. Briefly, 150 µl/well of the conditioned media were applied to the ELISA plate in duplicates. Fresh FMM media, which was not exposed to neurons and thus does not contain tau, was used as a background control. Each incubation step was performed for one hour at room temperature. Plates were read with the MESO QuickPlex SQ 120 (Meso Scale Discovery) after a 5 min incubation in 1X Read Buffer.

### Brain samples

Anonymized human post-mortem tissue was obtained from the London Neurodegenerative Diseases Brain Bank and the Southwest Dementia Brain Bank, members of the Brains for Dementia Research Network. Tissue donor characteristics were as follows: Control 1: male, age at death (AAD) 87, post-mortem interval (PMI) 48 h, cause of death (COD) cardiac failure. Control 2: male, AAD 77 years, PMI 16.75 h, COD unknown. Control 3: male, AAD 72 years, PMI 16.25 h, COD multiple myeloma. Control 4: female, AAD 96 years, PMI 72 h, COD urinary sepsis. Control 5: male, AAD 78 years, PMI 30.5 h, COD renal carcinoma. AD 1: female, AAD 83 years, PMI 70.75 h, COD endometrial carcinoma. AD2: female, AAD 95 years, PMI 87.5 h, COD frailty of old age. AD3: female, AAD 91 years, PMI 27.5 h, COD frailty of old age. AD4: male, AAD 80 years, PMI 50 h, COD pneumonia. AD5: male, AAD 81 years, PMI 38 h, COD Lewy body dementia.

The brain samples were homogenized with a Dounce homogenizers (Carl Roth) with Triton lysis buffer ((150 mM NaCl; 20 mM Tris, pH 7.5; 1 mM EDTA; 1 mM EGTA; 1% Triton-X-100; 1X Protease inhibitors; 1X Phosphatase inhibitors (both Roche), 500 µM IOX1, 2 µM Daminozide, 10 µM Trichostatin A, 5 mM Nicotinamide, 10 µM Paragyline hydrochloride, 1 µM Thiamet G) and placed into a 1.5 mL Eppendorf tube. Lysis was promoted by scraping the tubes over a hard surface and incubating the samples on ice for 10 min twice. Samples were centrifuged for 20 min at 12 000x g at 4°C, and the protein amount of the supernatant was determined via BCA assay (BioRad). For each sample two times 250 µg protein was used for immunoprecipitation (IP). For one IP, 250 µL Dynabeads® Protein G (Thermo Fisher) were incubated with a combination of 4 µg Tau12 (Biolegend, cat no 8065), 4 µg Tau 5 (Abcam, cat no ab80579) and 4 µg HT7 (Thermo Fisher, cat no MN1000) antibodies for 30 min at RT rotating. Then, 250 µg brain lysate was added for further incubation overnight at 4°C rotating. The beads were washed with lysis buffer before the tau was eluted with 250 µL elution buffer (50 mM glycine HCl, pH 2.8). Eluates were neutralized with 1 M Tris, pH 8-9, followed by a volume reduction to approx. 30 µL with Vivaspin columns (GE Healthcare). One tenth of the eluates was analyzed via WB to detect tau; while nine tenth of the samples were analyzed via Coomassie staining after separation on 7% NuPAGE™ Tris-Acetate protein gels (Thermo Fisher) in NuPAGE™ Tris-Acetate SDS running buffer (Thermo Fisher). The Western blot was stained with DAKO-Tau (1:10 000, Dako/Agilent, cat no A0024), while the second gel was stained with Colloidal Coomassie Staining solution (0.1% Coomassie Blue G250, 1M ammonium sulfate, 30% methanol, 3% o-phosphoric acid).

### Sample preparation for LC-MS/MS

Coomassie-stained bands were excised, chopped into small pieces and transferred to 0.5 ml Eppendorf tubes. For all following steps, buffers were exchanged by two consecutive 15 min incubation steps of the gel pieces with 200 µl of acetonitrile (ACN), whereby the ACN was removed after each step. Proteins were reduced by the addition of 200 µl of a 10 mM DTT solution in 100 mM ammonium bicarbonate (AmBiC, Sigma Aldrich, A6141) and incubation at 56°C for 30 min. Proteins were alkylated by the addition of 200 µl of a 55 mM chloroacetamide (CAA) in 100 mM AmBiC and incubation for 20 min in the dark. Samples were subjected to either an in-gel tryptic- or to an in-gel AspN digest. To this end, 0.1 µg/µl stock solutions of trypsin (Promega, V511A) or AspN (Promega, 90053) in resuspension buffer (Promega, V542A) were prepared and subsequently diluted with ice-cold 50 mM AmBiC buffer to achieve a final concentration of 1 ng/µl. Gel pieces were incubated in 50 µL of diluted enzyme for 30 min at 4°C, followed by overnight incubation at 37°C. Gel pieces were sonicated for 15 min, spun down and the supernatant was transferred into a glass vial (VDS Optilab, 93908556). Remaining gel pieces were washed with 50 µl of an aqueous solution of 50% ACN and 1% formic acid while sonicating for 15 min. The combined supernatants were dried down in a speedvac at 30°C. Peptides were reconstituted in 10 µl of an aqueous solution of 0.1% (v/v) formic acid.

### LC-MS/MS

Peptides were separated using nanoAcquity UPLC (Waters) with a nanoAcquity trapping (nanoAcquity Symmetry C18, 5µm, 180 µm × 20 mm) and analytical column (nanoAcquity BEH C18, 1.7µm, 75µm × 200mm), which was coupled to an LTQ Orbitrap Velos Pro (Thermo Fisher) using the Proxeon nanospray source. Peptides were loaded for 6 min using a constant flow of solvent A (0.1 % formic acid in H_2_O) at 5 µL/min. Peptides were then separated via the analytical column using a constant flow of 0.3 µl/min. Thereby, the percentage of solvent B (acetonitrile, 0.1 % formic acid) was increased from 3 to 10% within 5 min, followed by an increase to 40% within 10 min. Eluting peptides were ionized with a Pico-Tip Emitter 360 µm OD × 20 µm ID (10 µm tip, New Objective) applying a spray voltage of 2.2 kV at a transfer tube temperature of 300°C. Peptides were analyzed using data-dependent mode. Full scan MS spectra with a mass range of 300-2000 m/z were acquired in profile mode with a resolution of 30.000 and a filling time of 500 ms applying a limit of 1e6 ions. The 10 most intense ions were isolated (width 1.5 m/z) and fragmented in the HCD cell using a normalized collision energy of 30. Fragment masses were analyzed in the Orbitrap at a resolution of 7 500. 3e4 ions were selected within 150 ms and their fragmentation was achieved upon accumulation of selected precursor ions. MS/MS data were acquired in centroid mode of multiple charged (2+, 3+, 4+) precursor ions. The dynamic exclusion list was restricted to 500 entries with a maximum retention period of 30s and relative mass window of 10 ppm. In order to improve the mass accuracy, a lock mass correction using a background ion (m/z 445.12003) was applied.

### Data analysis

Acquired LC-MS/MS data were processed using IsobarQuant (doi:10.1038/nprot.2015.101) and Mascot (v2.2.07) using a reversed Uniprot homo sapiens database (UP000005640) including common contaminants. The following modifications were taken into account: Carbamidomethyl (C) (fixed modification), Acetyl (N-term) and Oxidation (M) (variable modifications). The mass error tolerance for full scan MS spectra was set to 10 ppm and for MS/MS spectra to 0.5 Da. A maximum of 2 missed cleavages were allowed. A minimum of 2 unique peptides with a peptide length of at least seven amino acids and a false discovery rate below 0.01 were required on the peptide and protein level. Only peptides corresponding to semi-trypsin or semi-AspN digests were considered.

Immunoblotting data were analyzed using Image Studio Lite (Li-cor Biosciences) and statistical analysis was performed with GraphPad Prism 7 (GraphPad Software) using the test noted within the respective figure legend.

## Results

Tau fragments truncated at N167 and N368 are present in control and AD human brain Tau fragmentation is an important part of the neuropathology observed in AD, and many tau fragments of unknown etiology have been previously observed in Western blot and mass spectrometry experiments (1, 2). In order to identify fragments that occur early in the disease process, we started our analysis of endogenous tau cleavage events with human entorhinal cortex samples from donors classified as either Braak stage 0-I (no tau pathology, hereafter referred to as control) or Braak III-IV, which corresponds to an early, often pre-symptomatic disease stage during the progression of AD (24). We furthermore decided to analyze a detergent-soluble fraction of the tissues, since we were most interested in proteolytic events occurring before the formation of mature tau tangles. We first enriched a Triton-X soluble brain extract for tau via immunoprecipitation with a combination of three monoclonal tau antibodies (Tau 12, HT7, Tau 5), which each target different epitopes and should thus enable us to target a broad range of fragmented tau and full-length tau species (Suppl. Fig. 1). We then applied two enzymatic digests in parallel, one with trypsin and one with AspN, as we found that these proteases yielded highly complementary peptides, leading to high sequence coverage when combined (overall coverage for all samples ranged from 73-91%, Suppl. Fig. S1). In the mass spectrometry data, we next searched specifically for tau peptides that contained either C- or N-termini that were not due to the proteolytic cleavage by trypsin or AspN digest, respectively. Such peptides thus represent physiological proteolytic events that occurred in the samples prior to their preparation for mass spectrometric analysis. With the requirements of a Mascot confidence score of ≥20 and the prerequisite that the respective site needed to be present in at least three separate brain samples, we identified a total of 19 cleavage sites. For more than half of these sites, we identified at least two peptides (Suppl. Table S1). Five sites have been reported previously (1, 12, 25, 26), while the remaining 14 represent potential new tau cleavage sites. For most of these sites, the corresponding proteases are unknown, but cleavage site prediction tools revealed potential proteolysis by a variety of enzymes including caspases, calpains, and MMPs (Suppl. Table S1).

The most prominent proteolysis event at N167 was detected with a total of four different peptides and was observed in nine out of ten brains when applying a cutoff ≥20 for the Mascot score (Suppl. Table S1 and Table 1). Two additional highly prevalent proteolysis events were P172 (9 brain samples, 3 peptides) and N368 (6 brain samples, 3 peptides) (Suppl. Table S1). Tau cleavage at both of these sites has been reported previously (12, 25). Interestingly, both aa167 and aa368 are asparagines, and N368 is a known cleavage site for asparagine endopeptidase (AEP) which has been associated with tau pathology in AD (12). We therefore decided to further investigate tau proteolysis at N167.

**Table 1:**
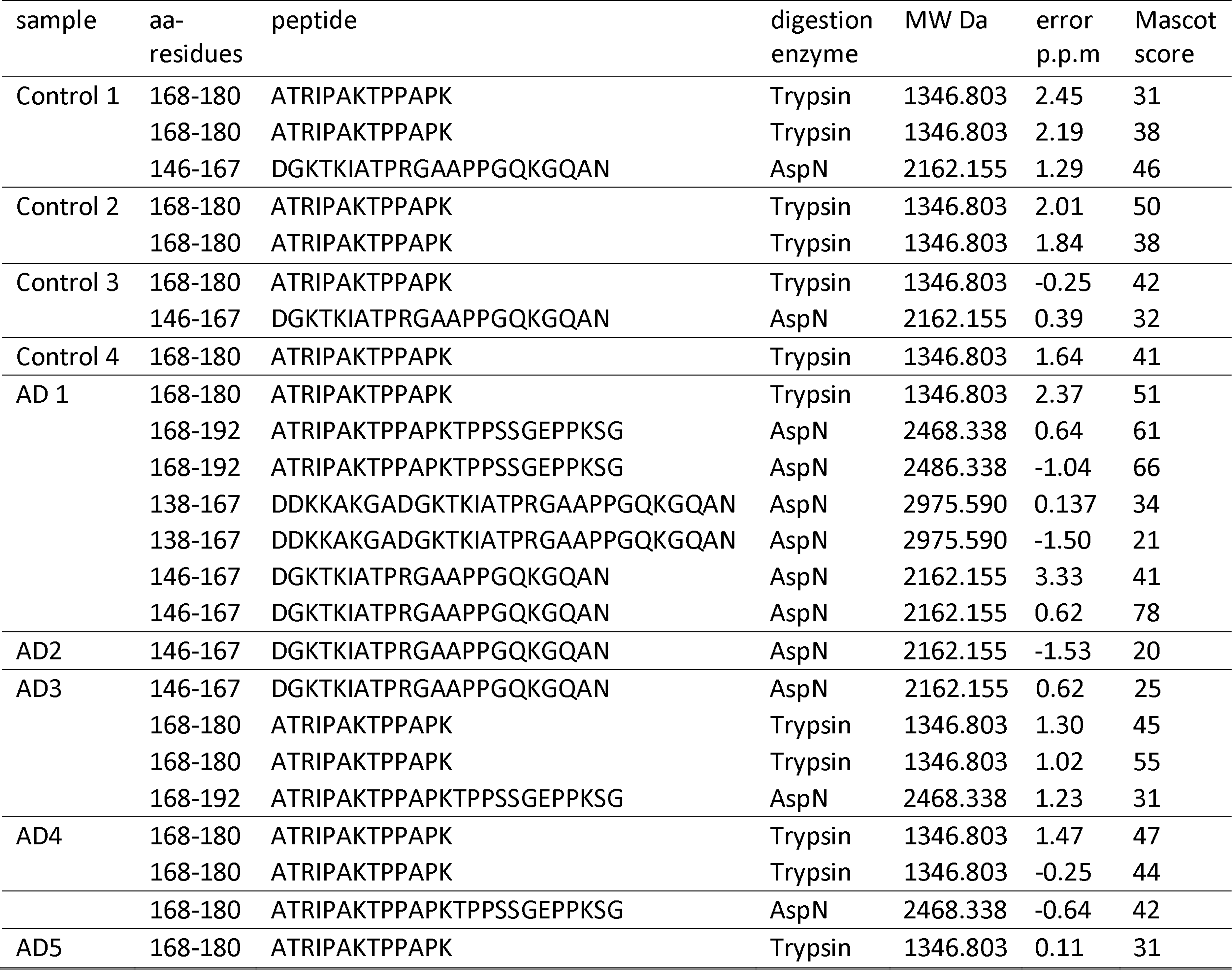
List of tau fragments truncated at N167 identified by mass spectrometry in control and AD human brain samples. Peptide sequences are numbered corresponding to the longest tau isoform (tau 2N4R, 441 aa).

### AEP cleaves tau at N167 and N368 in vitro

In order to verify that AEP is the enzyme responsible for tau cleavage at both N167 and N368, we next expressed either wildtype or cleavage-resistant tau mutants in combination with AEP in HEK-293 cells. We then evaluated the pattern of tau cleavage fragments detected with the antibodies Tau5 (epitope aa 210-241) and Tau N368, an antibody recognizing only tau fragments C-terminally truncated at N368 (12), as well as the C-terminal antibody DAKO-tau and an antibody recognizing the N-terminal HA tag on our tau constructs (Fig. 1A and B).

**Fig. 1:**
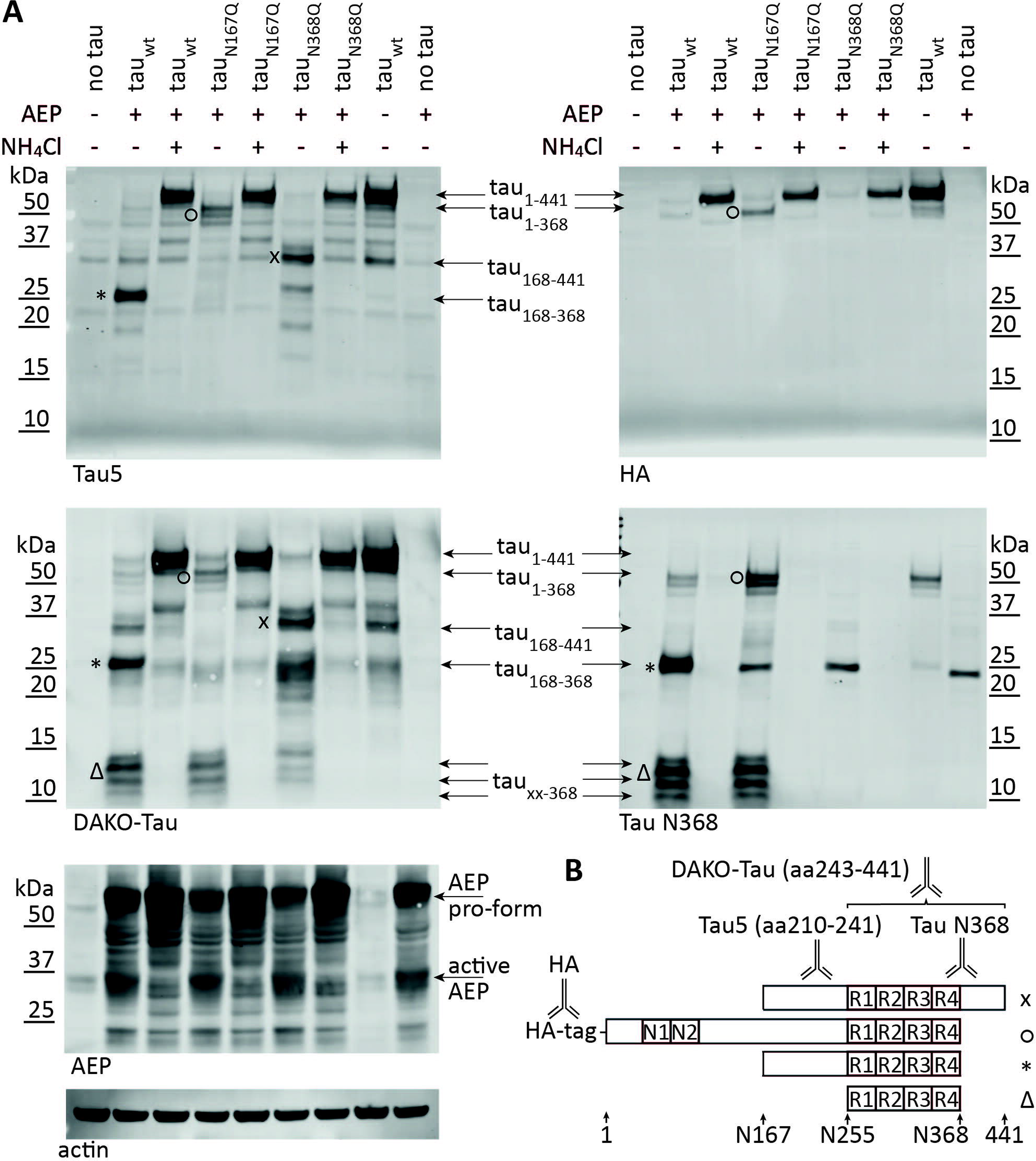
Tau is cleaved at N167 and N368 upon co-expression with AEP. **(A)** HEK293 cells were co-transfected with AEP and different tau constructs, and the tau cleavage pattern was analyzed by Western blotting with different antibodies (Tau5, DAKO-Tau, Tau N368 and HA). Major bands corresponded to cleavage events at N167 and N368, and are shifted or abolished when tau is mutated at these sites. 11: tau_1-368_, x: tau_168-441_, *: tau_168-368_, Δ: tau_xx-368_. Cells treated with NH_4_Cl show reduced tau cleavage. The expression and activation of AEP was validated in the same samples by Western blotting. Blots are representative of three independent experiments. **(B)** Graphic representation of the epitopes for Tau antibodies used in **(A)**.

Upon co-transfection with AEP, tau_wt_ is cleaved to a major fragment of 25 kDa, with almost no detectable full-length tau signal remaining (Fig. 1A, asterisk). This band is detected with the N368 antibody, Tau5 and DAKO-Tau, but does not contain the HA tag, suggesting it is truncated at N368 as well as at the N-terminus. While AEP transfection alone or in combination with tau mutants yields an additional band that is only detected with Tau N368, the 25 kDa band in the tau_wt_+AEP co-transfected sample is stronger and additionally positive for Tau5 and DAKO-tau (Fig. 1A), suggesting that the tau fragment overlaps with a non-specific band in the N368 blot.

We next compared the cleavage pattern of tau_wt_ with those obtained for tau protein mutated at the cleavage sites N167 and N368. The co-expression of both tau_N167Q_ and tau_N368Q_ with AEP abolished the cleavage fragment at 25 kDa, confirming its identity as tau_168-368_ (Fig. 1). In addition, tau_N167Q_ stabilizes bands at approx. 50 kDa that co-stain with Tau5, Tau N368 and DAKO-Tau as well as the HA tag and are likely processing intermediates truncated at N368, and one of which is likely tau_1-368_ (Fig. 1A, circle). In contrast, the tau_N368Q_ mutation prevents all staining with the Tau N368 antibody, as expected, and leads to a strong increase in a band of approx. 35 kDa that is detected with both Tau5 and DAKO-Tau (Fig. 1A, x). This band is absent in the tau_N167Q_ mutant and does not contain the HA tag at the N-terminus, therefore it likely corresponds to the tau_168-441_ fragment. A number of small bands around 10 kDa are detected only with the C-terminal DAKO-tau antibody and Tau N368 (Fig. 1A, triangle), suggesting that they terminate in N368 but have different N-terminal ends. One of these may be tau_256-368_, truncated at the previously described N255 AEP cleavage site (12), with an expected molecular weight of 12 kDa. Treatment with NH_4_Cl abolishes all differences in cleavage pattern between tau_wt_ and the tau mutants, confirming that these events are pH-dependent. As expected, the band for active AEP is also absent in NH_4_Cl treated samples, strongly suggesting that the pH-dependent activity of the overexpressed protease is responsible for the observed tau cleavage events.

### The tau_168-368_ fragment is aggregation prone

Since tau fragments are often more prone to aggregation than the full-length protein (1, 2, 12), we next determined the aggregation potential of the newly identified tau_168-368_. As commonly done for tau aggregation assays, we used heparin as an inducer and measured the increase in fluorescence caused by binding of the dye thioflavin T (ThT) to β-sheet rich protein aggregates (12). While tau_1-441_ did not aggregate under our conditions and in the timeframe of the experiment (Fig. 2A), the fluorescent signal for tau_168-368_ increased strongly 2 h after starting the reaction, and the maximum was reached between 5-10 h (Fig. 2A). While tau_1-441_ did not aggregate on its own under our conditions, we next investigated whether aggregation seeds derived from the tau168-368 fragment could promote the aggregation of full-length tau. To this end, we briefly sonicated pre-formed tau_168-368_ aggregates and added 5% as seeds to an aggregation reaction containing tau_1-441_. While the seed alone did not yield an appreciable fluorescence signal, we observed a time-dependent increase of fluorescence in the seeded tau_1-441_ aggregation reactions that reached a maximum after approximately 30 h of incubation (Fig. 2B). The addition of pre-formed aggregate seeds furthermore resulted in an aggregation kinetics with a very short lag time (Fig. 2B), in agreement with previous studies on seeded tau aggregation (27). This suggests that small amounts of aggregates formed by the tau_168-368_ fragment may cause full-length tau to form fibrils, even though the protein is normally much more conformationally stable and remains soluble.

**Fig. 2:**
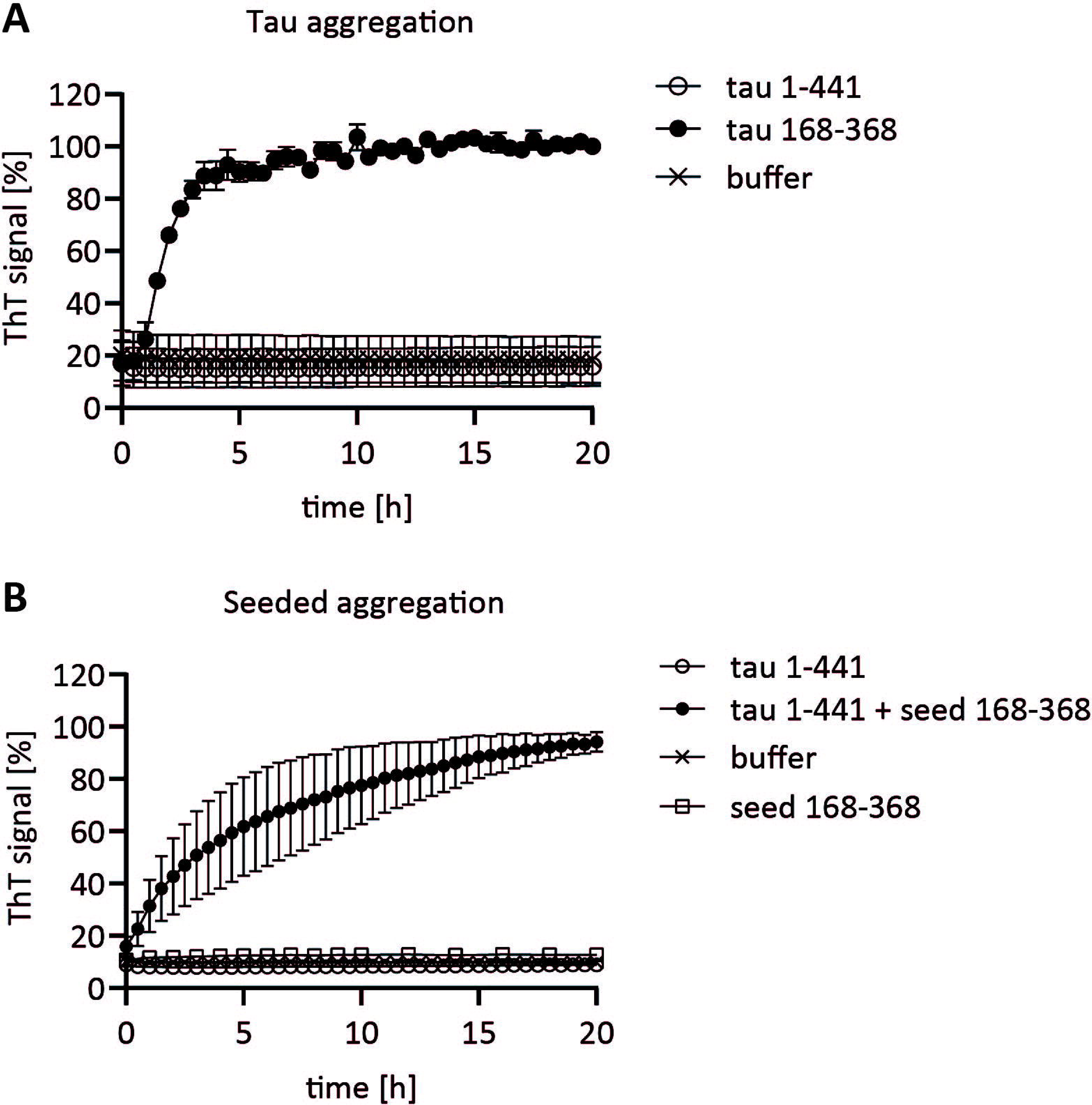
Tau_168-368_ is aggregation prone and efficiently seeds the aggregation of tau_1-441_. **(A)** The aggregation of recombinant tau_168-368_ was measured over time using the amyloid-binding dye thioflavin T (ThT). Full-length tau_1-441_ did not form ThT positive aggregates under the same conditions. Statistical significance was determined by two-way repeated measures ANOVA (time p < 0.0001, sample type p < 0.0001). Tukey’s post hoc analysis was used to determine that the tau_168-368_ sample is significantly different from both tau_1-441_ and buffer starting from the 2 h time point onwards. **(B)** The aggregation of full-length tau_1-441_ can be triggered through the addition of small amounts of pre-aggregated tau_168-368_ seeds. Statistical significance was determined by two-way repeated measures ANOVA (time p < 0.0001, sample type p < 0.0001). Tukey’s post hoc analysis was used to determine that the tau_1-441_+seed sample is significantly different from tau_1-441_, seed only and buffer starting from the 2 h time point onwards.

### AEP activity, but not the tau_168-368_ fragment is increased in AD patient brain samples

While our mass spectrometry study detected both N167 and N368 tau cleavage in brains from controls and AD patients alike, the method was not quantitative and thus not suitable to investigate potential disease-relevant differences in tau processing. Furthermore, mass spectrometry did not allow us to assess whether the tau_168-368_ fragment exists as such in human brain or if only single cleavage events were observed in vivo. We therefore analyzed our cohort of human brain tissues by Western blotting. As expected, Tau5 antibody staining yielded many bands between 75-20 kDa in size, corresponding to different tau isoforms and cleavage fragments (Fig. 3A). The Tau N368 antibody on the other hand stained a major band at 25 kDa, corresponding in size with the recombinant tau_168-368_ fragment (Fig. 3A), as well as a minor band at approx. 20 kDa. This band may correspond to the 3R isoform of the tau_168-368_ fragment, which lacks the R2 domain (Fig. 1B). Both bands also overlap with Tau5 staining. Quantification of the tau_168-368_ 4R fragment in a total of 9 control and 9 AD brain samples showed no difference in band intensity, suggesting that the fragment is present at equal levels in control and AD brains (Fig. 3B). Interestingly, Tau N368 did not stain any bands at the height expected for the tau_256-368_ fragment (approx. 12 kDa, as observed for the recombinant protein), in agreement with the fact that we did not detect the previously reported N255 cleavage (12) site by mass spectrometry, even though the area of the protein was well covered by peptides. We furthermore did not detect differences in the overall levels of tau in our samples as determined by quantification of the Tau5 signal (Fig. 3C).

**Fig. 3:**
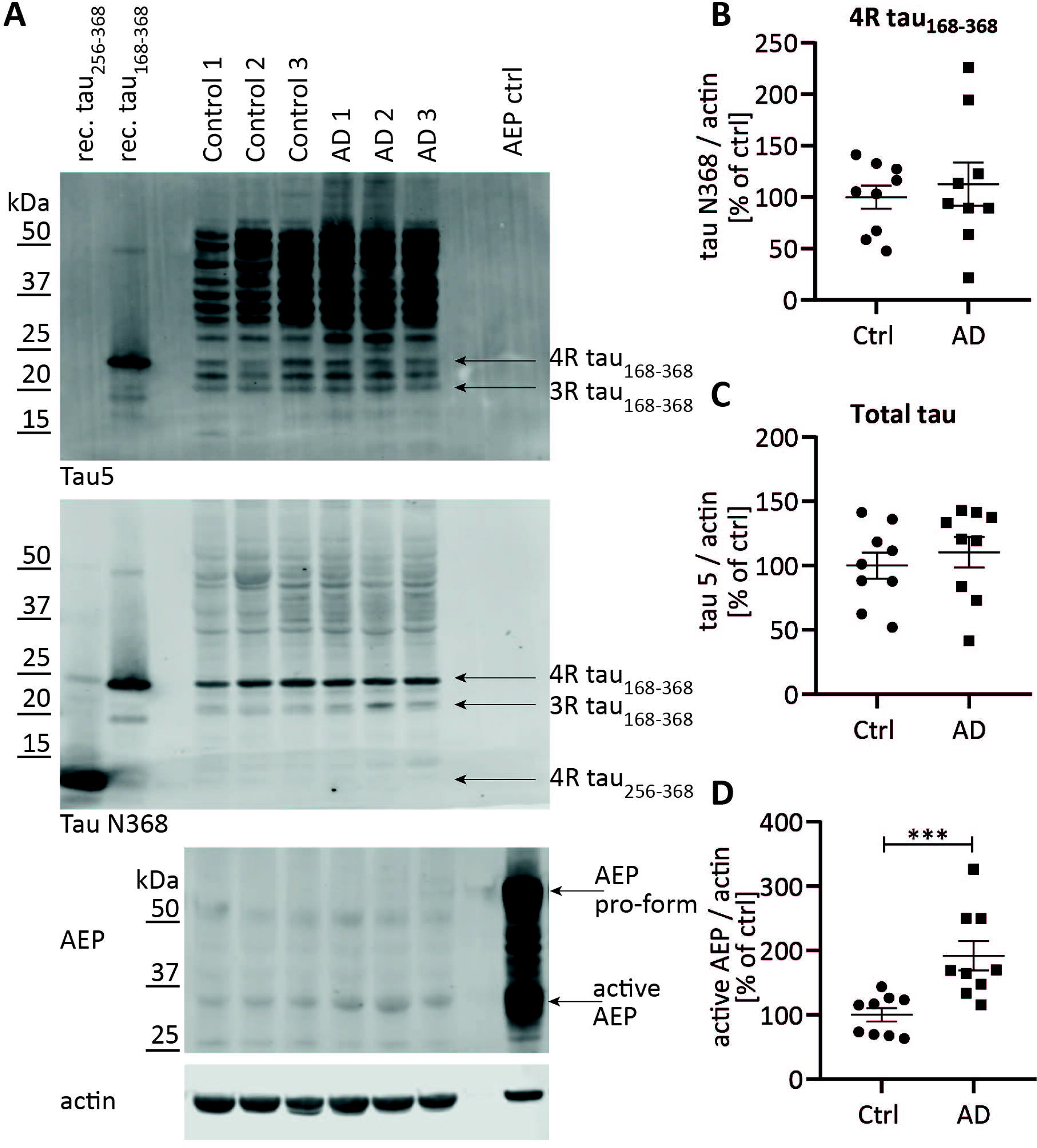
Analysis of tau fragmentation and AEP activation in human brain samples. **(A)** Representative Western blots demonstrating tau isoforms and cleavage fragments detected with the total tau antibody Tau5. Staining with Tau N368 reveals a major band at the size corresponding to the 4R tau168-368 fragment, and a minor band that may represent the same fragment for the 3R tau isoform. AEP is detected in its active form at approx. 35 kDa. **(B) – (D)** Quantifications of the Western blots shown in **(A)** reveal no difference in the levels of the tau_168-368_ fragment **(B)** or total tau (**(C)**, all bands in the lane quantified together) between control and AD patient brain tissues, while active AEP is significantly increased in AD samples **(D)**. Statistical significance was determined by Mann Whitney test, ***: p < 0.001.

Previous studies have suggested that AEP activity increases both with aging and in AD (12, 16). We therefore checked for the presence of the active AEP fragment in control and AD brain tissues and found a significant increase in our cohort of 9 AD patient samples compared to controls (Fig. 3D). While this seems to be contradictory to the unaltered levels of the tau168-368 fragment, it is possible that AEP-mediated tau cleavage is a physiological event occurring in control as well as AD patient brains and may only be an intermediate step in the proteolytic degradation of tau without any accumulation of tau_168-368_.

### Full-length tau is cleaved upon uptake into AEP-expressing cells

AEP is a predominantly lysosomal enzyme and is active at an acidic pH (13). Since tau can be secreted and taken up by cells, and the described mechanisms for tau uptake deliver their cargo to endo/lysosomes, we were wondering if AEP-mediated cleavage of tau may occur during this process. We started by adding recombinant, monomeric tau_1-441_ or the tau_168-368_ fragment to the medium of HEK-293 cells, which do not produce endogenous tau. When we subsequently analyzed cell lysates for the presence of tau, we observed that both tau_1-441_ and the tau_168-368_ fragment were readily taken up; however, we did not observe any cleavage (Fig. 4A). This may be due to non-detectable amounts of endogenous AEP in HEK-293 cells (Fig. 4A). In contrast, when we transfected HEK-293 cells with AEP, tau_1-441_ was taken up and cleaved to tau_168-368_, as observed by staining with the Tau5 and Tau N368 antibodies (Fig. 4A). NH_4_Cl treatment slightly decreases the amount of tau_168-368_ while increasing the detectable level of full-length tau, paralleled by decreased levels of active AEP (Fig. 4A). Interestingly, we did not observe any further processing of tau_168-368_ after uptake (not shown), and the levels of this fragment also did not change upon AEP transfection or NH_4_Cl treatment (Fig. 4A). In tau_1-441_ treated cells we observed an additional band at approx. 35 kDa, which was positive for Tau5 staining, but negative for Tau N368 (Fig. 4A), and we therefore concluded that it likely corresponds to the tau_168-441_ processing intermediate (Fig. 1A). As observed previously (Fig. 1A), AEP transfection alone leads to a non-specific band at approx. 25 kDa detected by Tau N368, which overlaps with the tau_168-368_ signal (Fig. 4A).

**Fig. 4:**
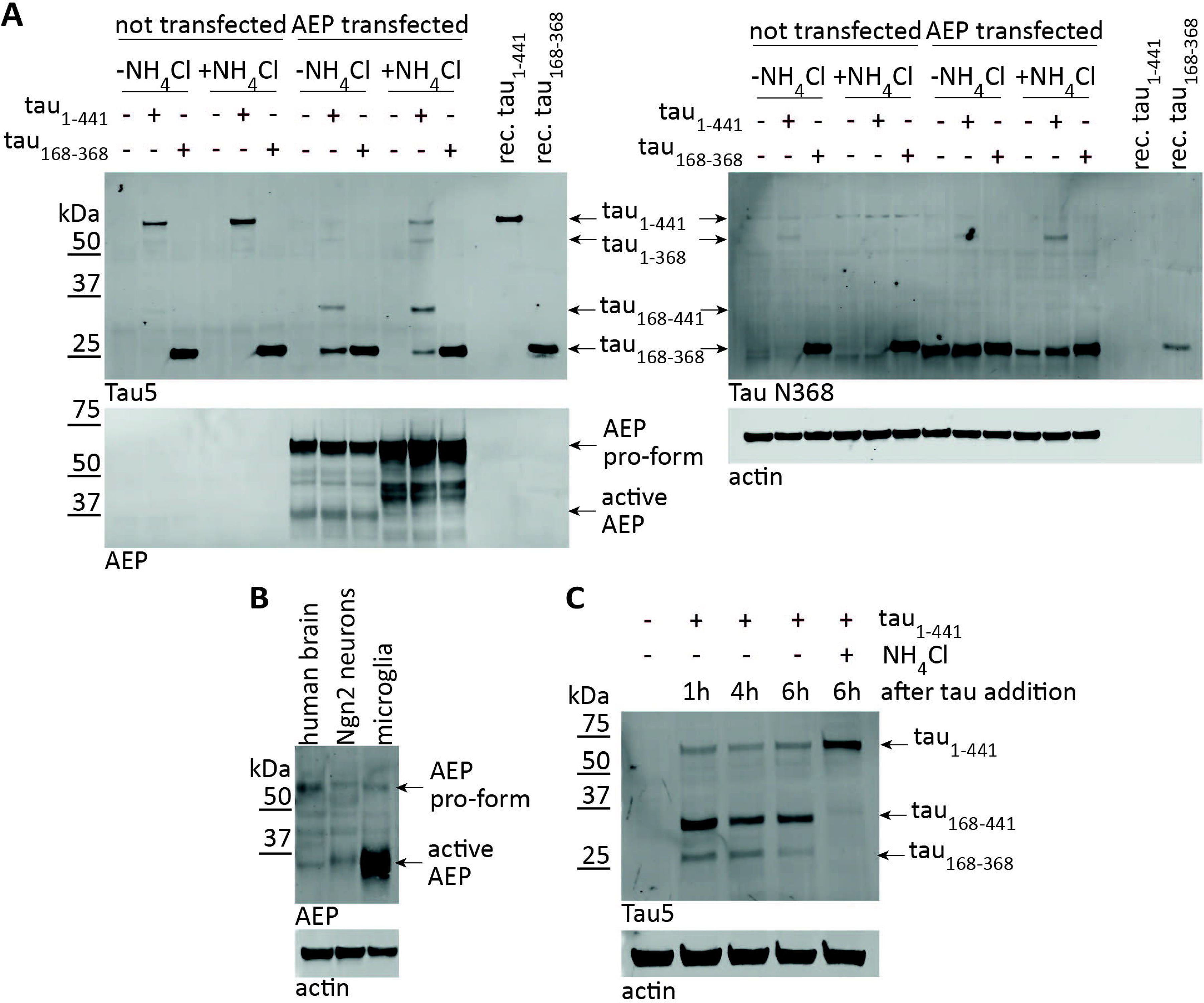
Recombinant tau is endocytosed and cleaved upon uptake into AEP-expressing cells. **(A)** Representative Western blots demonstrating the uptake of recombinant tau_1-441_ and ^tau^168-368 in HEK293 cells. Cleavage of endocytosed tau is observed, if the cells overexpress AEP. The tau fragments generated correspond to tau_1-368_, tau_168-441_ and tau_168-368_, and cleavage is reduced in the presence of NH_4_Cl. **(B)** Representative Western blot demonstrating strong expression of active AEP in a microglia cell line, but not in iPS-derived neurons or total brain extracts. **(C)** Representative Western blot demonstrating the uptake of recombinant tau_1-441_ in microglia cells. Cleavage of endocytosed tau is observed in the absence of NH_4_Cl only, but the fragments do not accumulate over time.

### Microglia, but not neurons express sufficient AEP to cleave tau

Our results thus far show that the presence of active AEP is a prerequisite for tau cleavage at N167 and N368, and previous studies suggest that AEP is highly expressed in antigen-processing cells (15). In agreement with these data, we found high levels of active AEP in microglia, the primary phagocytes in the brain (Fig. 4B). Neurons derived from induced pluripotent stem cells (iPS) on the other hand exhibited very low levels of AEP (Fig. 4B). We then assessed which of these two cell types was able to cleave tau after uptake from the culture medium. Since neurons express large amounts of tau protein endogenously, we used recombinant, biotinylated tau_1-441_ protein for this assay. Western blot analysis of neuronal lysates using a streptavidin antibody demonstrated that exogenous tau was internalized, but specific cleavage fragments sensitive to NH_4_Cl treatment were not detected (Suppl. Fig. S2). Microglia, on the other hand, took up tau_1-441_ and digested it to a main fragment of approx. 35 kDa, a size corresponding to tau168-441 (Fig. 4C). Interestingly, this fragment did not accumulate over time, while cleavage was prevented by NH_4_Cl treatment, confirming the pH-dependent activity of the protease (Fig. 4C).

To investigate whether tau is completely removed by microglia, we next assessed microglia medium 24 h after the addition of recombinant tau_1-441_. While full-length tau was readily detected in medium incubated in the absence of cells, the presence of microglia led to the complete disappearance of tau protein during this time period (Fig. 5A). Tau removal was not affected by the addition of NH_4_Cl, suggesting that tau uptake can occur even during lysosomal dysfunction. Our data furthermore suggest that microglia degrade the endocytosed protein rather than secrete tau fragments after the initial proteolysis step, since no re-released fragments were detected.

**Fig. 5:**
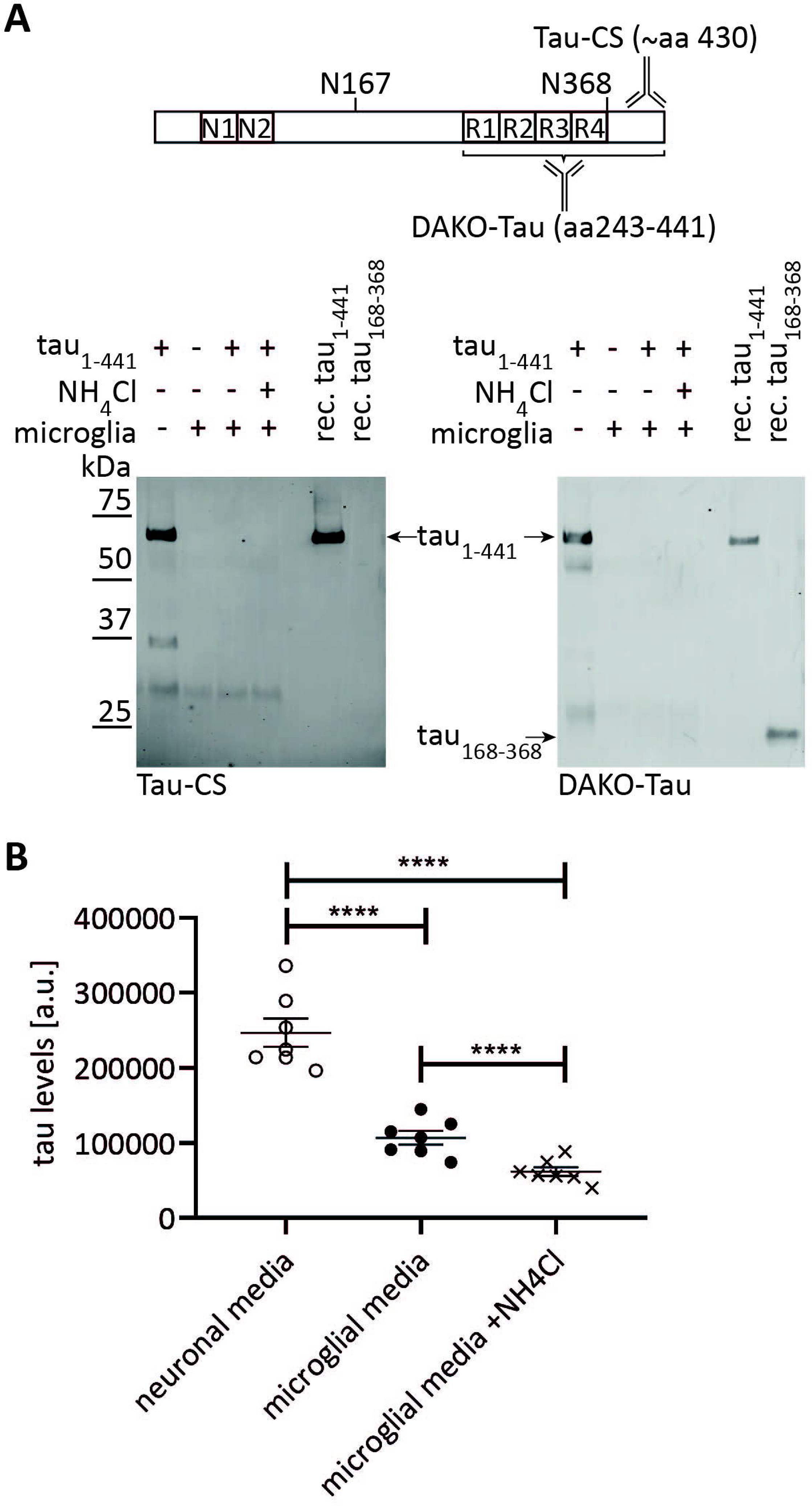
Microglia completely remove recombinant tau from the medium and also process tau released from iPS-derived neurons. **(A)** Microglia were incubated with recombinant tau_1-441_ in the medium for 24 h, as a control, recombinant tau was incubated in cell culture medium without microglia present. Media was then subjected to immunoprecipitation with Tau5 antibody. Representative Western blots with both DAKO-Tau and Tau-CS demonstrate that no full length or fragmented tau remains in the medium when microglia are present, and NH_4_Cl treatment did not prevent tau removal. **(B)** Medium derived from day 15 iPS-derived neurons was transplanted onto microglia, and tau levels were determined by electrochemiluminescence ELISA after incubation for 24 h. The addition of NH_4_Cl during the last 6 h of incubation further reduced tau levels. Statistical significance was determined by repeated measures one-way ANOVA (p < 0.0001) and post-hoc Holm-Sidak’s test. ****: p < 0.0001.

### Microglia remove tau previously secreted by neurons

In order for microglia to take up and digest tau in the brain, the protein first needs to be secreted by neurons. Several studies have shown that tau secretion is a physiological process, possibly regulated by neuronal activity (4, 5, 28). Since neuronal tau may differ from the recombinant protein in its tertiary structure or due to post-translational modifications, we next assessed the ability of microglia to remove tau protein secreted from neuronal cultures. Conditioned medium was obtained from iPS cells differentiated into neurons through the inducible expression of Ngn2 (23). These cells exhibit a neuronal morphology and robustly express tau protein starting at day 15 after differentiation (Suppl. Fig. S3), and we also detected tau released into the culture medium at this timepoint using a sensitive electrochemiluminescence ELISA method (Fig. 5B). To investigate the effect of microglia on the secreted tau protein, we transplanted conditioned neuronal medium onto microglial cells. After an incubation period of 24 h we found that the tau levels in the medium had significantly decreased (Fig. 5B), similar to the effect previously observed for recombinant tau (Fig. 5A). Interestingly, the addition of NH_4_Cl during the last 6 h of incubation led to an even stronger decrease of tau levels (Fig. 5B). This confirms that microglia can efficiently take up tau even in the absence of functional lysosomal cleavage and degradation pathways, and the increased tau uptake is furthermore in agreement with a previous report showing that microglial endocytosis is facilitated by NH_4_Cl-mediated changes in intracellular pH (29).

## Discussion

In this study, we used a mass spectrometry approach to identify tau cleavage events in human brain tissue from AD patients and age-matched control subjects. We identified a previously unknown, but highly prevalent proteolytic event at N167 of tau, which was detected with high confidence in nine out of ten brain samples analyzed. In vitro studies revealed that this site is processed by AEP, a protease that has previously been linked to AD and was shown to cleave tau at both N368 and N255 (12). While we confirmed processing at the N368 site in our samples, we did not observe tau cleavage at N255 in human brain, although our mass spectrometric approach revealed peptides mapping this region. However, it is possible that N255 is only processed subsequent to other cleavage events, generating fragments that would have migrated on the SDS-PAGE below the area we excised for their mass spectrometric analysis. This is consistent with our in vitro findings, where the only bands that were consistently generated by AEP from tau_wt_, tau_N167Q_ and tau_N368Q_ (and may thus correspond to N255 cleavage) were small (10-15 kDa). Taken together this suggests that the site can be processed downstream of other cleavage events, but fragments may not be very abundant in vivo.

Previous studies have shown that AEP activity is increased both during aging and in AD brain tissue (12, 16). We also found a significant increase in active AEP enzyme in tissues from AD patients compared to controls, however this did not result in an increase in the tau_168-368_ fragment. In fact, the equal abundance of tau_168-368_ in control and AD patient brains suggests that tau cleavage by AEP is a physiological event, and the lack of tau fragment accumulation over time in vitro furthermore points towards tau_168-368_ being an intermediate product of a degradation pathway.

AEP has been described as an important protease in inflammatory signaling, and one of its unique substrates is TLR9 (15, 30). Accordingly, we also found that microglia, the resident macrophage cells of the brain, express more AEP than neurons and mostly contain the active form of the enzyme. While aberrant AEP activity in neurons has been reported under special conditions (14, 31), we do not observe AEP-mediated tau cleavage in iPS-derived neurons. Even after endocytotic uptake of recombinant tau, which delivers tau directly to endolysosomes and should thus make it available for AEP cleavage (4, 10), we do not observe appreciable amounts of tau_168-368_ fragments, suggesting that this proteolytic event is not primarily taking place in neurons.

On the other hand, cells that do contain active AEP such as transfected HEK cells or microglia, readily cleave tau after uptake from the medium, generating fragments truncated at N167 and N368. The cleavage process, but not the uptake is inhibited by the lysosomal acidification inhibitor NH_4_Cl. In microglia, tau uptake eventually leads to the complete removal of all tau protein from the culture medium without an intracellular accumulation of tau fragments, suggesting that the protein is not only cleaved by AEP but completely broken down and removed. However, our experiments do not exclude the possibility that small amounts of pathogenic forms of tau may escape from microglia and contribute to the spreading of pathology in vivo, which may be promoted by microglial dysfunction in the disease state (32, 33). In fact, our in vitro experiments suggest that tau_168-368_ is aggregation prone and, once misfolded, can seed the aggregation of full-length tau. Nevertheless, tau processing in microglia can only have an effect on spreading and pathology if the protein is not fully degraded after uptake, for example due to endolysosomal dysfunction (11), post-translational modifications or misfolding events that prevent tau cleavage (34). Therapeutic approaches targeting such processes may therefore also remove any disease-associated blockage in tau degradation without potentially interfering with a physiological tau cleavage event.

## Conclusions

Previous studies have suggested that AEP may be a therapeutic target in several neurodegenerative diseases, due to its increased activity and capability to generate potentially toxic fragments from different disease-related proteins such as tau, APP or α-synuclein (12, 16, 17, 35). However, our results suggest that tau proteolysis by AEP is a physiological event that occurs as a part of normal tau degradation, and the inhibition of this process may in fact be harmful, especially since AEP is also an important player in TLR-mediated immune signaling (15). Since AEP activity in the brain mostly localizes to microglia, the increased activity of the enzyme observed in AD may reflect neuroinflammatory phenotypes commonly observed in the disease and may thus be ameliorated by approaches targeting aberrant microglia activation rather than AEP itself (36).

## Abbreviations

aa: amino acid
AAD: age at death
AD: Alzheimer’s disease
AEP: asparagine endopeptidase
APP: amyloid precursor protein
approx.: approximately
COD: cause of death
ctrl: control
ELISA: Enzyme-linked immunosorbent assay
fl: full-length
HEK: human embryonic kidney cells
iPSC: induced pluripotent stem cell
kDa: kilodalton
MW: molecular weight
Ngn2: Neurogenin 2
PMI: post-mortem interval
p.p.m.: parts per million
ThT: Thioflavin T
wt: wild-type

## Acknowledgements

The authors thank Christian Weber for assistance with the culture of iPS-derived neurons, and Dr. Kim Remans and Dr. Jacob Scheurich at the EMBL protein purification core facility for the expression and purification of recombinant tau proteins. We are indebted to Dr. Martin Fuhrmann and Dr. Laura Gasparini for fruitful discussions and advice and Dr. Theron Johnson for access to the Meso Scale Discovery Quickplex platform. Human post-mortem tissue was obtained from the London Neurodegenerative Diseases Brain Bank and the Southwest Dementia Brain Bank, members of the Brains for Dementia Research Network.

## Availability of data and materials

The datasets used and/or analysed during the current study are available from the corresponding author on reasonable request.

## Authors’ contributions

AB performed cell culture and cell-free experiments, analyzed data, coordinated coauthor contributions and prepared the first draft of the manuscript. MB performed and analyzed experiments with human brain samples, EEH performed and analyzed ELISA experiments, PH performed mass spectrometry experiments and analyzed the resulting data, DCS, MW and SAF performed iPS culture and differentiation, EM performed site-directed mutagenesis. DEE supervised the project, performed experiments with microglia, analyzed data and wrote the manuscript. All authors read and approved the final manuscript.

## Ethics approval and consent to participate

Human brain samples were collected with informed consent by the London Neurodegenerative Diseases Brain Bank and the Southwest Dementia Brain Bank and were provided in a strictly anonymized fashion. hiPSCs were derived from fibroblasts that are part of the NIA Aging Cell Repository at the Coriell Institute for Medical Research (37). Informed consent was obtained by Coriell and the fibroblasts were provided in a strictly anonymized fashion for iPS derivation.

