## Supplementary figures and images for "Asparagine endopeptidase cleaves tau at N167 after uptake into microglia"

### Suppl. Fig. S1

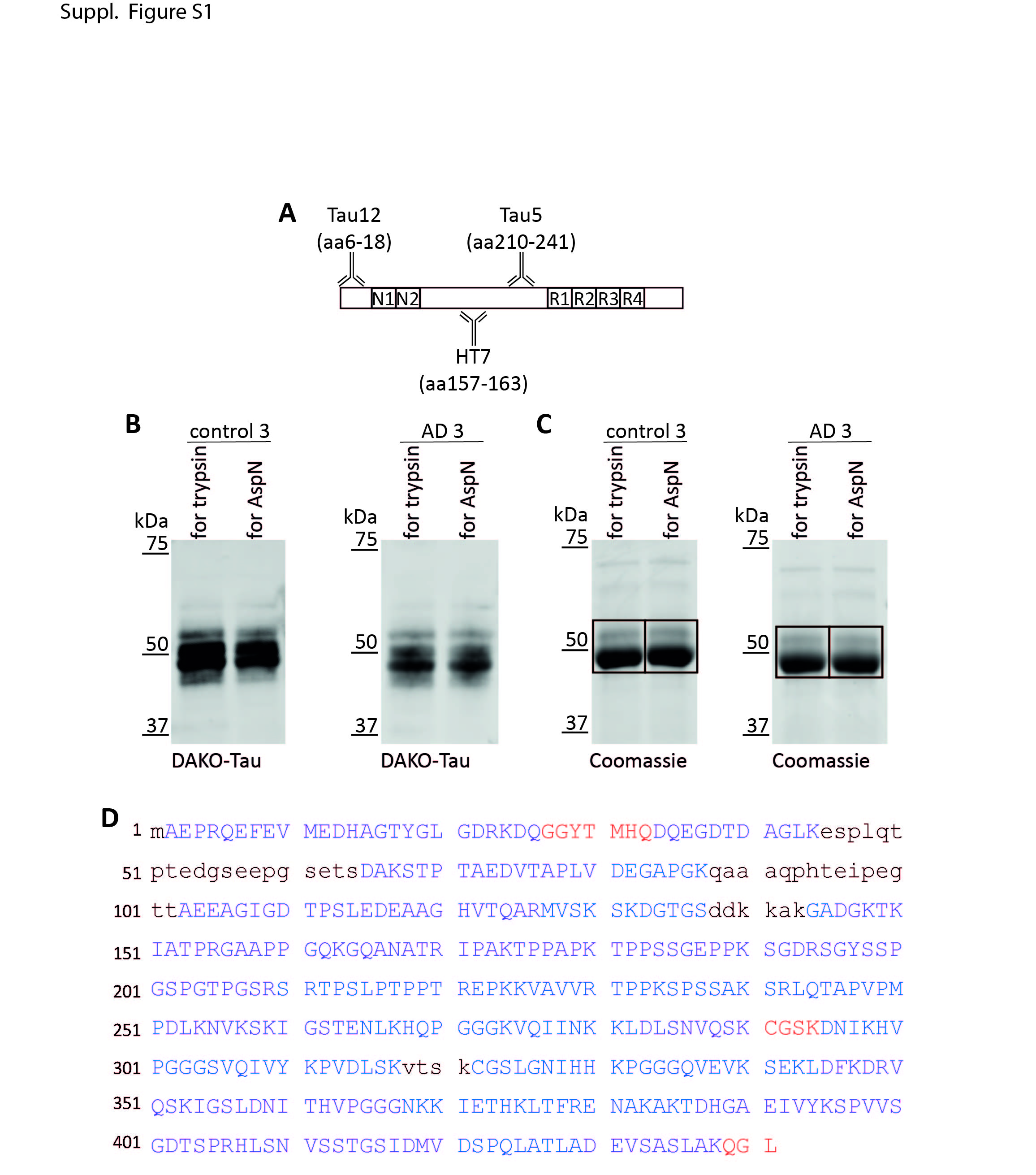

### Suppl. Fig. S2

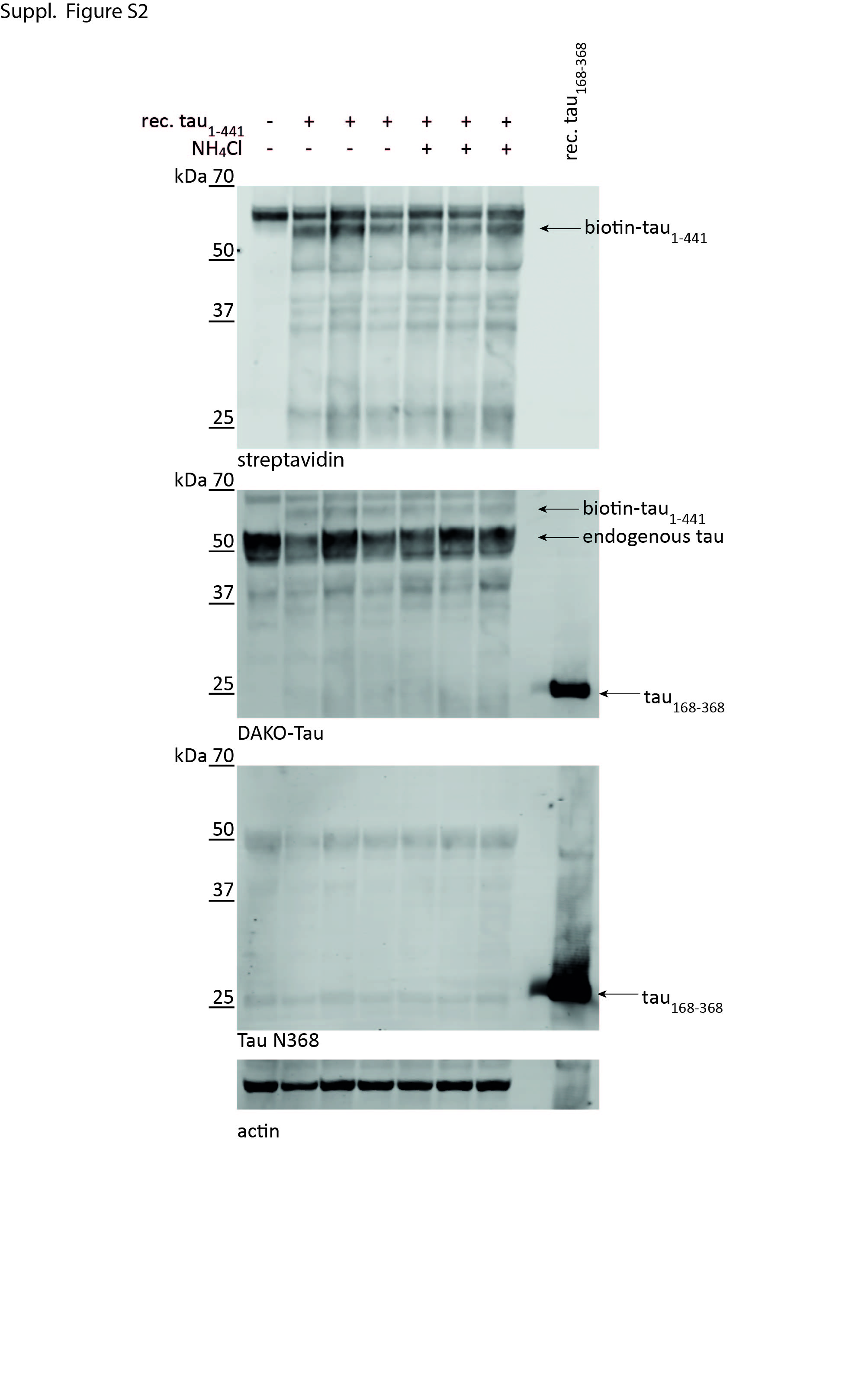

### Suppl. Fig. S3

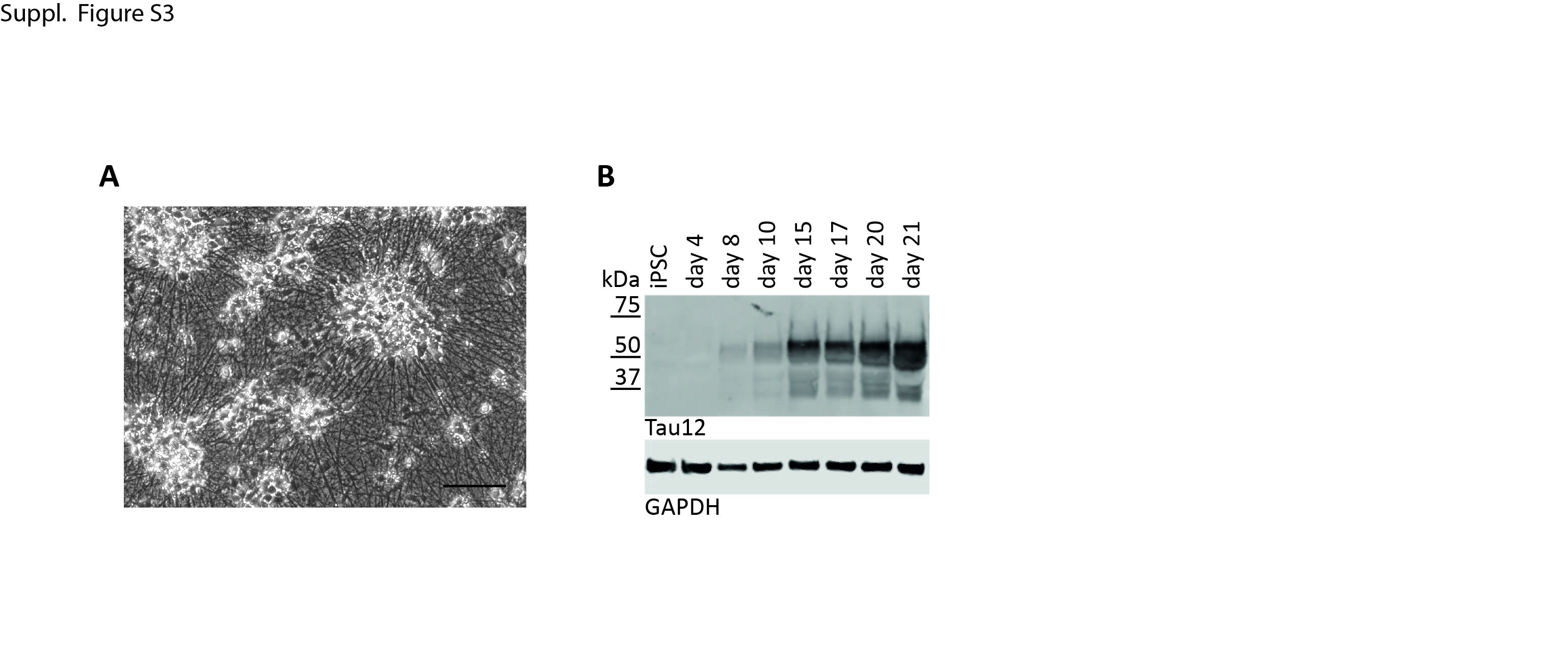
