## Supplementary material for "Asparagine endopeptidase cleaves tau at N167 after uptake into microglia": Suppl. Figure legends

**Supplemental Figure legends:**

**Suppl. Fig. S1: Band excision strategy and sequence coverage obtained by mass spectrometry.**

**(A)** Epitopes of the monoclonal antibodies used for immunoprecipitation of tau from human brain samples. **(B)** Representative Western blots and **(C)** Coomassie-stained gels demonstrating major tau bands observed after immunoprecipitation from human brain samples. The boxed areas were excised and subjected to mass spectrometry analysis. **(D)** Sequence coverage obtained for a representative brain sample (AD3). Purple: sequence covered by both trypsin and AspN-derived peptides. Red: sequence covered by AspN-derived peptides. Blue: sequence covered by trypsin-derived peptides.

**Suppl. Fig. S2: Recombinant tau is endocytosed, but not cleaved by AEP in iPS-derived neurons.**

Biotinylated, recombinant tau was taken up into neurons and detected by streptavidin-coupled antibody staining. While both tau_1-441_ and cleavage fragments were observed, no bands at the appropriate size for tau_168-368_ (25 kDa) were detected, and none of the fragments was sensitive to NH_4_Cl treatment, suggesting that pH-dependent proteases such as AEP do not contribute to the observed cleavage.

**Suppl. Fig. S3: Morphology and tau expression patterns in iPS-derived neurons.**

**(A)** iPS-derived cells display neuronal morphology at day 15 of the differentiation process. Scale bar: 100 µm. **(B)** iPS-derived neurons express tau protein from day 15 onwards.
