## Supplemental Data 1 for "Asparagine endopeptidase cleaves tau at N167 after uptake into microglia"

| Brain sample(s) | Enzyme | Peptide(s) | Cleavage site | Reference |
| --- | --- | --- | --- | --- |
| Control 1, Control 3, Control 5, AD 1, AD 2, AD 3, AD 4 |  | EDHAGTYGLG | M11 | (1, 2) |
| Control 1, Control 2, Control 3, Control 5, AD 1, AD 3, AD 4 |  | GAAPPGQKGQANATRIPAKTPPAPKTPPSSGEPPKSG | R155 |  |
| Control 1, Control 2, Control 3, Control 5, AD 1, AD 2, AD 3, AD 4, AD 5 | AEP | ATRIPAKTPPAPK  DGKTKIATPRGAAPPGQKGQAN  ATRIPAKTPPAPKTPPSSGEPPKSG  DDKKAKGADGKTKIATPRGAAPPGQKGQAN | N167 |  |
| Control 1, Control 2, Control 3, Control 5, AD 1, AD 3, AD 4, AD 5 | MMP-2, MMP-3, MMP-12, KLK-5, PCSK2 | RIPAKTPPAPKTPPSSGEPPKSG | R170 |  |
| Control 1, Control 2, Control 3, Control 4, Control 5, AD 1, AD 4, AD 5 |  | GQANATRIP  GAAPPGQKGQANATRIP  AKTPPAPKTPPSSGEPPKSG | P172 | (1, 2) |
| Control 3, AD 1, AD 5 | MMP-3, MMP-7, PCSK2 | LQTAPVPMP | R242 |  |
| Control 3, AD 1, AD 5 |  | GKVQIINKKL  IGSTENLKHQPGG | G272 |  |
| Control 1, Control 2, Control 3, Control 4, AD 1, AD 3, AD 4 | MMP-3, MMP-9, neprilysin | IGSTENLKHQPGGGKVQ | Q276 |  |
| Control 1, Control 2, Control 3, Control 5, AD 3, AD 4 |  | IGSLDNITHV | V363 |  |
| Control 2, Control 3, AD 3 |  | IGSLDNITHVPG  IGSLDNITHVPGG | G366 |  |
| Control 2, Control 3, AD 1, AD 2, AD 3, AD 4 | AEP | DNITHVPGGGN  IGSLDNITHVPGGGN  DRVQSKIGSLDNITHVPGGGN | N368 | (2, 3) |
| Control 1, Control 3, Control 4, AD 1, AD 3, AD 4 | MMP-3, KLK-4, KLK-5 | ENAKAKTDHGAEIVYKSPVVSG | R379 |  |

| Brain sample(s) | Enzyme | Peptide(s) | Cleavage site | Reference |
| --- | --- | --- | --- | --- |
| Control 3, Control 4, AD 1, AD 4 | MMP-3, MMP-7, neprilysin, granzyme B | IVYKSPVVSG | E391 | (1, 2) |
| AD 2, AD 3, AD4, AD 5 | calpain | TDHGAEIVY  AKTDHGAEIVY | Y394 |  |
| Control 3, Control 4, AD 2 |  | AKTDHGAEIVYKSPVVS  GDTSPRHLSNVSSTGSI  GDTSPRHLSNVSSTGSIDMV | S400 |  |
| Control 1, Control 2, Control 3, Control 5, AD 1, AD 3, AD 4, AD 5 | calpain | TDHGAEIVYKSPVVSGDT  AKTDHGAEIVYKSPVVSGDT | T403 |  |
| Control 1, Control 2, Control 3, Control 4, Control 5, AD 1, AD 3, AD 4, AD 5 | caspase | HLSNVSSTGSID | D418 |  |
| Control 1, Control 2, Control 3, AD 1, AD 3 | caspase | HLSNVSSTGSIDMVD  DTSPRHLSNVSSTGSIDMVD | D421 | (2, 4) |
| Control 2, AD 1, AD 3 | calpain | ASLAKQGL  HLSNVSSTGSIDMVDSPQLATLADEVS | S433 |  |

**Suppl. Table S1: Tau cleavage sites detected by mass spectrometry.**

Endogenous tau cleavage sites were extracted from semi-tryptic and semi-AspN peptides detected by mass spectrometry. All events with Mascot scores >20 that were detected in three or more individual brain samples are listed. Enzymes listed in grey were predicted using the following online tools: CaspDB (caspase cleavage prediction (5)), CaMPDB (calpain cleavage prediction (6)), iProt-Sub (other proteases (7)).
